# Neuronal adenosine A_2A_ receptors signal ergogenic effects of caffeine

**DOI:** 10.1101/2020.04.02.021923

**Authors:** Aderbal S Aguiar, Ana Elisa Speck, Paula M. Canas, Rodrigo A. Cunha

## Abstract

Ergogenic aid is a substance or method used for enhancing exercise and sports performance. Caffeine is the most used ergogenic aid for athletes, but the mechanisms are still unknown. Forty-two adult female (19±0.6 g) and 40 male mice (24±0.4 g) from a global and forebrain A_2A_R knockout and colony (FMUC, University of Coimbra) underwent an open field and ergospirometry exercise test. Caffeine (15 mg/kg, i.p.) and SCH 58261 (1 mg/kg, i.p.) were administered 15 minutes before the animals ran to exhaustion. We also evaluate the estrous cycle and infrared temperature (rest and recovery). Caffeine was psychostimulant in wild type females and males, but we observed this expected effect of SCH-58261 only in males. Caffeine and SCH-58261 were also ergogenic for wild type animals, that is, they increased running power and maximal O_2_ consumption (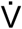O_2_max). The psychostimulant and ergogenic effects of caffeine and SCH-58261 disappeared in A_2A_R knockout females (global) and males (forebrain). The estrous cycle did not influence any evaluated parameters, as well as exercise-induced hyperthermia was similar between savages and knockouts. Our results suggest that the neuronal A_2A_R receptors signal the ergogenic effects of caffeine in female and male mice.

## INTRODUCTION

Caffeine is the most used ergogenic substance for athletes. Caffeine spike exercise performance in rodents^1-5^ and humans^6-8^. It increases endurance in submaximal cycling in humans^6-8^ and on the treadmill in rodents^1-5^. Candidate mechanisms are controversial due to *in vivo* toxicity, such as blocking GABA_A_ receptors, inhibition of phosphodiesterase and increased calcium mobilization achieved with millimolar concentrations of caffeine^9-10^. This is the evidence used by the sports sciences. However, adenosine receptors have a high caffeine affinity for caffeine^9-10^. We hypothesized that adenosine A_2A_ receptors (A_2A_R) are essential for the ergogenic effects of caffeine, and evaluated its effects in an exercise test.

## RESULTS

### Caffeine and SCH-58261 are not ergogenic in global A_2A_R knockouts

Ergospirometry evaluates exercise submaximal and maximal performance during ergometer exercise (power, O_2_ and CO_2_ kinetics) and exclude subjective measures of fatigue^1,3,11^. The test assessed the ergogenic effects of Caffeine and SCH-58261 (a potent and selective A_2A_R antagonist) and A_2A_R knockout (KO) mice. We used females in the first set of experiments, due to the characteristic of our animal colony. There is no impairment in the use of females for performance evaluation^1^. In the open field, the basal motor behavior was not different between wildtype (WT) and A_2A_R KO (t_18_ = 0.8, Fig.1A). SCH-58261 did not modify the locomotion (t_18_ = 0.4, Fig.1A), and caffeine was psychostimulant for WT animals, but not for A_2A_R KO mice (F_1,36_ = 5.8, P < 0.05, Fig.1A).

**Fig. 1.**
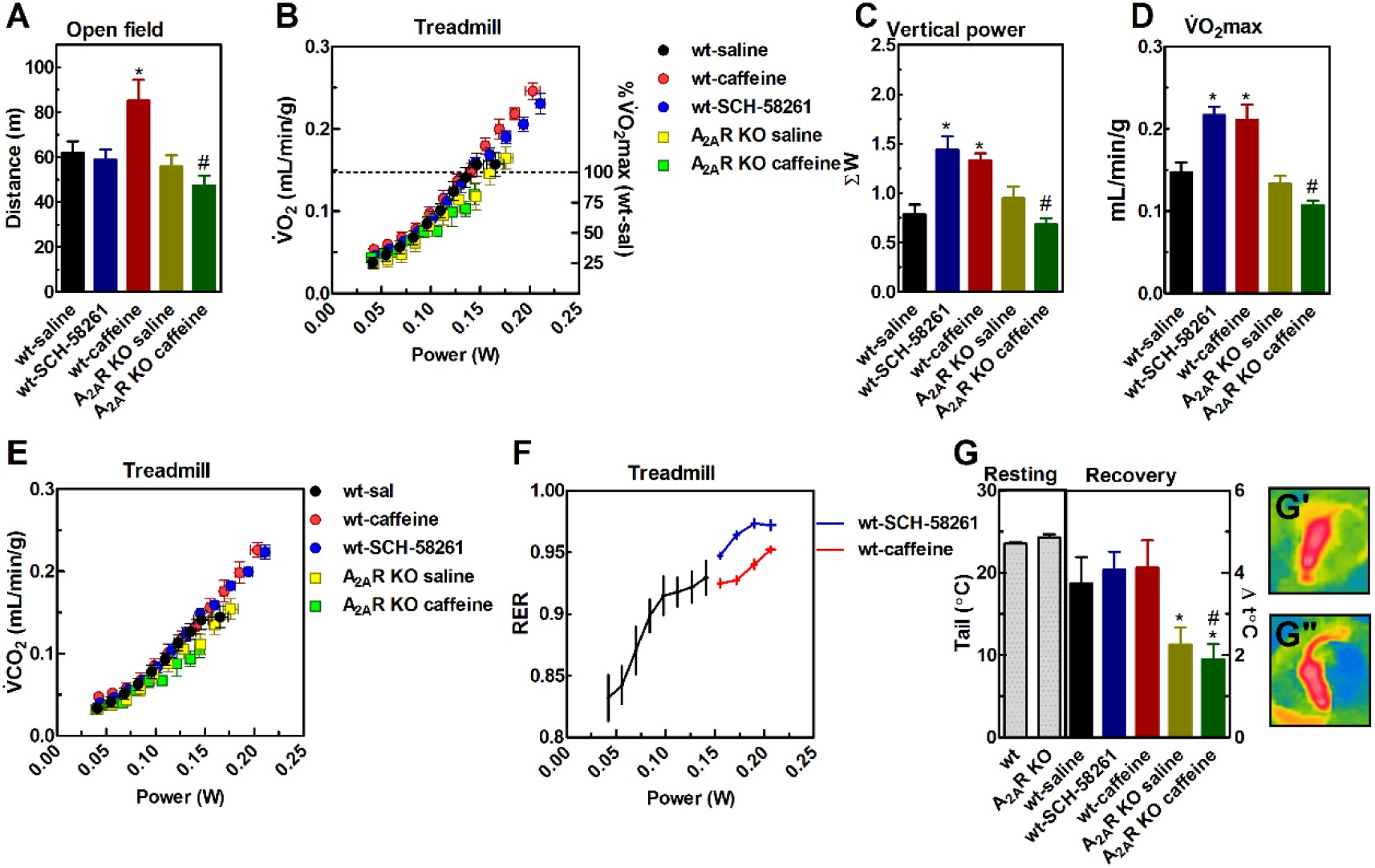
Effect of caffeine (15 mg/kg, i.p.) and SCH-58261 (1 mg/kg, i.p.) on the basal locomotion and exercise performance of female mice (A). Caffeine was psychostimulant in the open field, but not in caffeine-treated A_2A_R KO mice. Ergospirometry increased 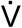O_2_ (B), 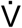CO_2_ (E), RER (D) and running power (B) until the animals reached fatigue. Caffeine and SCH-58261 increased running power (C) and 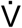O_2_max of WT (B), but not A_2A_R KO. Resting tail temperature was not different (G). Exercise-induced tail hyperthermia was greater in WT than in A_2A_R KO, with no caffeine or SCH-58261 effect. Infrared Fig.G’-G’’ Illustrates the tails lighting up with the heat. Values are expressed as mean ± standard error of the mean (SEM). N = 8-9 animals/group for 12 independent experiments. * P < 0.05 vs. saline for SCH-58261 (Student’s t-test) or caffeine (ANOVA, Bonferroni post hoc test). # P < 0.05 vs. wt-caffeine (ANOVA, Bonferroni post hoc test).

Ergospirometry increased running power (F_7,210_ = 6243, P < 0.05, Fig.1B) and O_2_ consumption (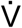O_2_, F_7,196_ = 255, P < 0.05, Fig.1B) in a gradual manner, without submaximal 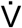O_2_ differences at speeds 15 → 50 cm/s (SCH-58261 F_7,77_ = 0.8; caffeine × A_2A_R KO F_7,133_ = 2.1). These speeds (15 → 50 cm/s) correspond to exercise test stages fulfilled by all groups, and also the maximal O_2_ uptake (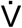O_2_max) of WT controls (wt-saline, Fig.1B – dotted line indicating 100%). High-performance speeds (55 → 65 cm/s) were mainly reached by WT animals treated with SCH-58261 or caffeine, reflecting the ergogenic profile of A_2A_R antagonists.

We demonstrated for the first time that SCH-58261 was ergogenic, that is, it increase running power (t_16_ = 2.1, P < 0.05, Fig.1D) and 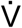O_2_max (t_16_ = 3.3, P < 0.05, Fig.1D) of WT females. The ergogenic effects of caffeine were demonstrated in rodents^2-6^. These effects were reproduced in our Lab by demonstrating increased running power (F_1,32_ = 21, P < 0.05, Fig.1C) and 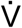O_2_max (F_1,32_ = 11, P < 0.05, Fig.1D). According to hypothesis, caffeine was not ergogenic for A_2A_R KO mice; caffeine unchanged running power (F_1,32_ = 21, P < 0.05, Fig.1C) and 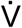O_2_max (F_1,32_ = 11, P < 0.05, Fig.1D).

Ergospirometry also evaluates substrate utilization during exercise through respiratory-exchange ratio (RER = 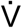CO_2_/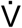O_2_). CO_2_ production (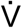CO_2_) had similar kinetics to 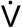O_2_ during the exercise test at speeds 15 → 50 cm/s (F_7,210_ = 257, P < 0.05, Fig.1E). RER increased during exercise (F_1,189_ = 15.8, P < 0.05, Fig.1F) at these speeds, without any effect of SCH-58261 (F_7,77_ = 0.8) or caffeine × genotype (F_7,133_ = 1.2).

Exercise and substrate oxidation produce heat; we have described tail hyperthermia as an index of exercise-induced thermoregulation^1^. That away, 3 females at estrous (Fig.S1C) were excluded due to large exercise-induced tail hyperthermia at this stage of estrous cycle^1^. The following results refer to females in diestrus (Fig.S1A), proestrus (Fig.S1B) and metestrus (Fig.S1D). Tail resting IR temperature was similar among wildtype and A_2A_R KO mice (t_34_ = 3.4, Fig.1G). The Fig.1G’ shows the IR profile of female tails at rest, and the tail heating (candle effect) caused by high-intensity exercise (Fig.1G’’). Tail temperature increased approximately 4±0.3°C for WT and 2.1±0.3°C for A_2A_R KO mice, with a significant effect of genotype factor (F_1,27_ = 15, P < 0.05, Fig.1G). There were no significant effects of SCH-58261 (t_16_ = 0.5, Fig.1G) or caffeine (F_1,27_ = 1.6, Fig.1G) on exercise-induced tail heating.

### The ergogenic effects of caffeine and SCH-58261 depend on neuronal A_2A_R receptors

These results suggest the robust role of A_2A_R in the ergogenic effects of caffeine. We use forebrain neuron-specific A_2A_R KO mice (fb-A_2A_R KO) to understand whether the role of A_2A_R is cell-type specific. ‘Floxed” A_2A_R mice were crossed with calmodulin-dependent protein kinase II α subunit (CaMKIIα)-Cre transgenic line^7^. This experiment was carried out with males, again due to the characteristics of our animal colony.

SCH-58261 (t_21_ = 2.3, P < 0.05, Fig.2A) and caffeine (F_1,34_ = 8.6, P < 0.05, Fig.2A) were psychostimulants for WT mice in the open field. But caffeine did not change the locomotion of fb-A_2A_R KO mice. There were no submaximal differences in ergospirometry for speeds 15 → 55 cm/s (Fig.2B-F). Fig.2C-D shows the ergogenic effect of SCH-58261 on running power (t_16_ = 3.3, P < 0.05) and 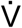O_2_max (t_16_ = 2.1, P < 0.05) in WT mice, as did caffeine (Power F_1,26_ = 10.7; 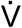O_2_max F_1,30_ = 5.0). Caffeine was no longer ergogenic in fb-A_2A_R KO mice.

**Fig. 2.**
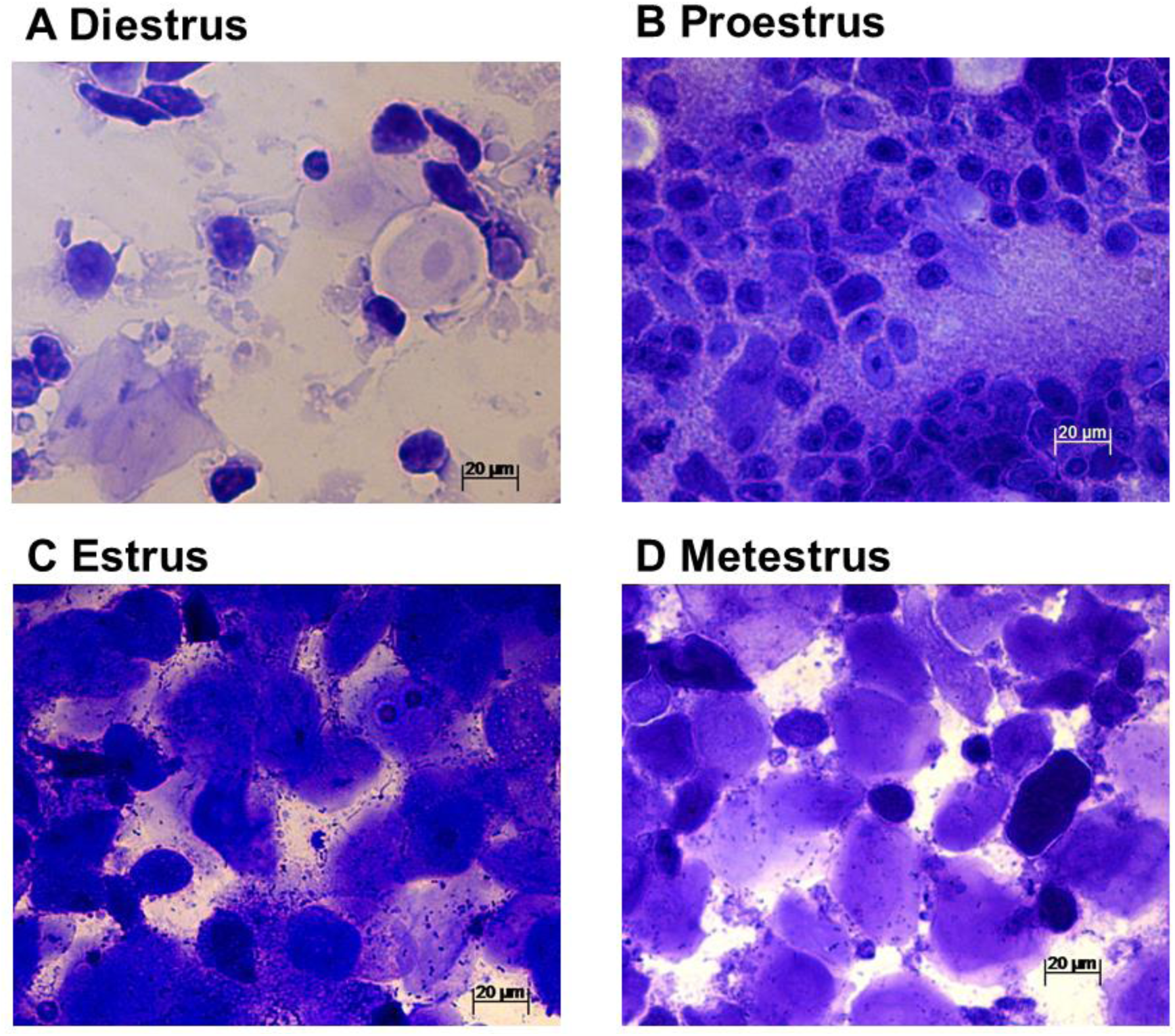
Effect of caffeine (15 mg/kg, i.p.) and SCH-58261 (1 mg/kg, i.p.) on the basal locomotion and exercise performance of male mice (A). Caffeine and SCH-58261 were psychostimulant in the open field, but not in caffeine-treated forebrain-A_2A_R KO mice. Ergospirometry increased 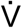O_2_ (B), 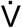CO_2_ (E), RER (D) and running power (B) until the animals reached fatigue. Caffeine and SCH-58261 increased running power (C) and 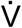O_2_max of WT (B), but not forebrain-A_2A_R KO. Tail temperature was not different at resting and after exercise (G). Values are expressed as mean ± standard error of the mean (SEM). N = 8-9 animals/group for 12 independent experiments. * P < 0.05 vs. saline for SCH-58261 (Student’s t-test) or caffeine (ANOVA, Bonferroni post hoc test). # P < 0.05 vs. wt-caffeine (ANOVA, Bonferroni post hoc test).

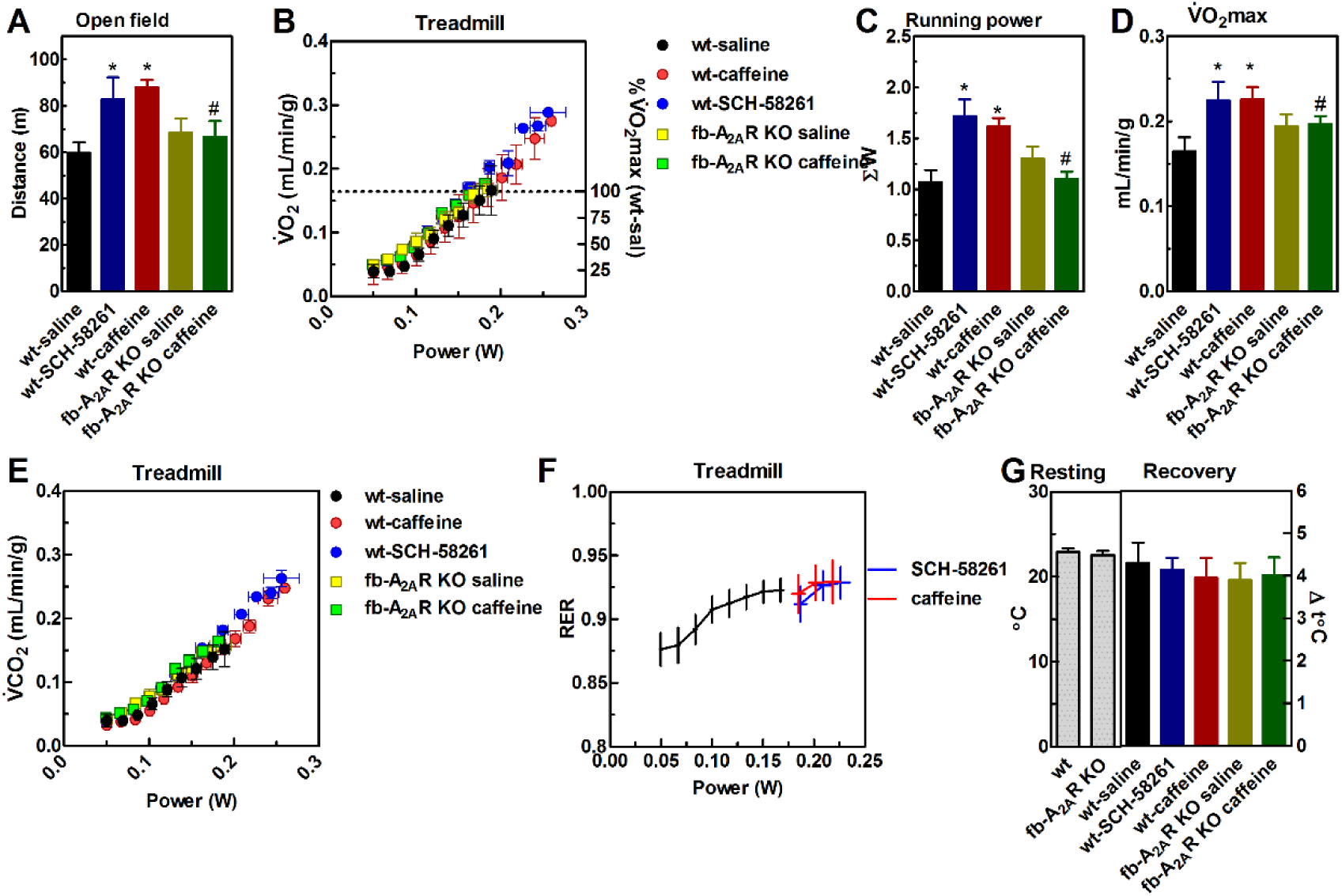

Then we see no differences in RER (Fig.2F) and tail temperature (Fig.2G). SCH-58261 (F_1,56_ = 0.3) and caffeine × genotype (F_1,98_ = 0.2) unchanged RER (Fig.2F). Tail temperature was similar at rest (t_30_ = 0.5) and after maximal exercise (F_1,26_ = 0.07) for all groups.

## DISCUSSION

Caffeine increases exercise performance in rodents^1-5^ and humans^6-8^. Early evidence demonstrated metabolic changes during exercise, through glycolysis/glycogenolysis inhibition (and glycogen economy) by increasing fat oxidation (and increased RER and thermogenesis)^6-7^. Authors speculated on the phosphodiesterase inhibition and calcium mobilization evidence^9-10^. But there is a serious lack of support for all these metabolic changes^4,13-17^. Increased fat oxidation should decrease RER. However, we did not observe submaximal differences in ergospirometry (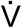O_2_, 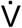CO_2_, and RER), only in peak exercise performance (power and 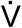O_2_max) in SCH-58261 and caffeine-treated WT animals. These effects were shown to be A_2A_R-dependent. The main substrate typically changed during the exercise test, from initially fat (RER ≈ 0.7) to carbohydrate (RER ≈ 1.0). This pattern remained even in A_2A_R KO mice. Caffeine and SCH-58261 also did not change exercise-induced tail heating, or thermogenesis. This effect of caffeine was been ruled out in exercising subjects^8,9^. Our data does not support the metabolic mechanisms of caffeine for its ergogenic effects.

Caffeine is lipophilic, easily crosses blood-brain barrier and reaches CNS 20-50 μM levels after 1-3 cups of coffee^9^, sufficient to antagonize A_1_R and A_2A_R. Caffeine is psychostimulant^9,19,20^ and decreases the rate of perceived exertion (RPE) during exercise^5,8,17^, which opens the central fatigue hypothesis of caffeine also for its ergogenic effects. Here, caffeine was not psychostimulant and ergogenic in fb-A_2A_R KO mice. There are two other pharmacological studies in this same line of evidence. The intracerebroventricular injection of caffeine (200 μg) was ergogenic in rats^4^, and the nonselective adenosine receptor agonist, 5′-(N-Ethylcarboxamido)adenosine (NECA) defeat this effect^4^. In a reverse experimental design, Zheng & Hasegawa^2^ showed that systemic caffeine reversed the poor running performance of NECA-treated rats. Highlight for the non-selective mechanism of NECA, in contrast to highly selectivity of SCH-58261 by A_2A_R. Our data reinforce the ergogenic role of A_2A_R antagonism in the CNS, which shows similarities to psychostimulant effects of caffeine and SCH-58261. The psychostimulant effects of caffeine depend on dopaminergic signaling. As for its less investigated ergogenic effect, caffeine increases extracellular levels of dopamine in the cerebrospinal fluid of running rats^2^. Caffeine is also a negative allosteric modulator of D_2_ receptors of A_2A_R-D_2_R G-protein coupled receptor (GPCR) heteromers in the basal ganglia^10,11^. Selective A_2A_R antagonists improve dopaminergic signaling through increased availability of D_2_/D_3_ receptors^11^, causing wakefulness and psychostimulation. Caffeine, methylphenidate (dopamine and norepinephrine reuptake inhibitor) and reboxetine (selective noradrenaline reuptake inhibitor) are ergogenic^12-14^ and reduce saccade eye fatigue after exercise, a marker of central fatigue^14,15^. Caffeine also reduces central fatigue induced by transcranial magnetic stimulation (TMS)^16^. In summary, our data supports the well-set role of neuronal A_2A_R for ergogenic effects of caffeine.

## METHODS

### Animal colony and drugs

The animals used were male (23.9 ± 0.4 g, 8-10 weeks old) and female mice (20 ± 0.2 g, 8-10 weeks old) from our global-A_2A_R knock-out (KO) mice (A_2A_R KO)^17^ and forebrain-A_2A_R KO (fb-A_2A_R KO)^18^ inbred colony and wildtype (WT) littermates. Cre-LoxP method generate fb-A_2A_R KO^18^. The generation and genotyping of A_2A_R KO and fb-A_2A_R KO mice were previously described^17,18^.

Mice were housed in HEPA-filtered ventilated racks and collective cages (n = 3-5) under controlled environment (12 h light-dark cycle, lights on at 7:00 AM, and room temperature of 21 ± 1°C) with *ad libitum* access to food and water. Housing and handling were performed according to European Union guidelines. The study was approved by the Ethical Committee of the Center for Neuroscience and Cell Biology (University of Coimbra).

Experimental design is shown in Fig.S2. The animals were habituated to handling, injections (0.9% NaCl – saline, i.p.) and treadmill for 3 days. SCH-58261 (1 mg/kg, i.p., dissolved in 10% DMSO in 0.9% NaCl – saline) and caffeine (15 mg/kg, i.p., dissolved in saline) were freshly prepared and administered systemically (volume 10 ml/kg body weight) on days 4 and 5, 15 minutes before open field (4^th^ day) and ergospirometry (5^th^ day). All behavioral tests and drug treatments were carried out between 9:00 and 17:00 hours in a sound-attenuated and temperature/humidity controlled room (20.3±0.6 °C, 62.8±0.4 %H_2_O) under low-intensity light (≈ 10 lux). Open field and treadmill were cleaned with 10% EtOH between individual experiments. Allocation to the experimental groups was random. For each test, the experimental unit was an individual animal.

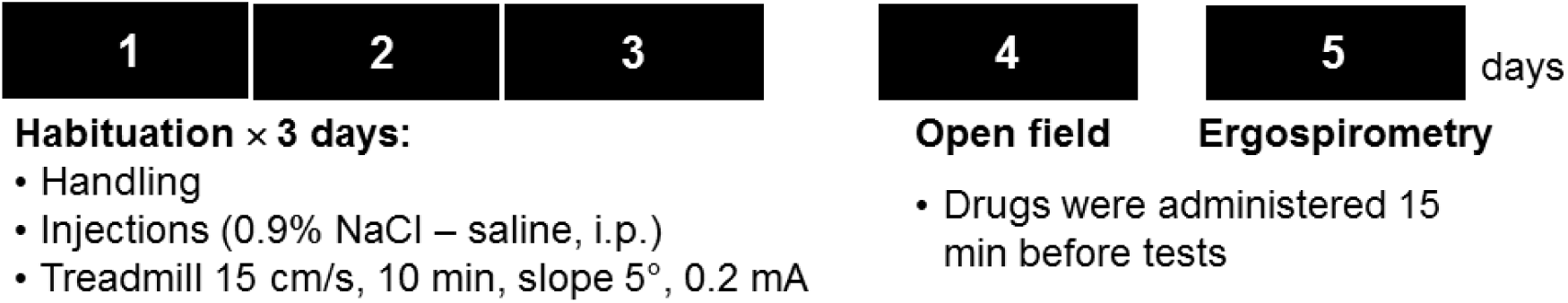

### Open field

The exploration of an open field (38 × 38 cm) was analyzed for 15 min using the ANY-maze™ video tracking system (Stoelting Co.).

### Ergospirometry

Mice were accustomed with a single-lane treadmill (Panlab LE8710, Harvard apparatus) at speed 15 cm/s (10 min, slope 5°, 0.2 mA) with 24 h interval between each habituation session (Fig.S2). The incremental running protocol started at 15 cm/s with an increment of 5 cm/s every 2 min at 5° inclination. The exercise lasted until running exhaustion, defined by the inability of the animal to leave the electrical grid for 5 seconds^1,19,20^. We estimated the power output for treadmill running based on a standard conversion of the vertical work, body weight and running speed^1,21,22^.

Oxygen uptake (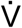O_2_) and carbon dioxide production (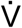CO_2_) were estimated in a metabolic chamber (Gas Analyzer ML206, 23 × 5 × 5 cm, AD Instruments, Harvard) coupled to treadmill. The animals remained in the chamber for 15 min prior to exercise testing. Atmospheric air (≈21% O_2_, ≈0.03% CO_2_) was renewed at a rate 120 mL/min, using the same sampling rate for the LASER oxygen sensor (Oxigraf X2004, resolution 0.01%) and infrared carbon dioxide sensor (Servomex Model 15050, resolution 0.1%).

### Vaginal cytology

We evaluated the estrous cycle immediately after the exercise test, through 4-5 consecutive vaginal lavages (with 40-50 μL of distillated H_2_O) then mounted on gelatinized slides (76 × 26 mm)^23,24^. These procedures lasted no more than 3-5 minutes, and there were no major temporal delays between behavioral experiments and fluid collection for vaginal cytology^1^.

The vaginal smear were desiccated at room temperature and covered with 0.1% crystal violet for 1 min, then twice washed with 1 mL H_2_O and desiccated at room temperature. The slides were mounted with Eukitt medium (Sigma-Aldrich) and evaluated under an optical microscope at 1x, 5x and 20x (Zeiss Axio Imager 2). The characterization of the estrous cycle was performed according to literature^23,24^. Females were categorized for initial (metestrus) or late (diestrus) follicular phase, ovulation (proestrus), or luteal phase (estrus)

### Thermal imaging

An infrared (IR) camera (FLiR C2, emissivity of 0.95, FLiR Systems) placed overtop (25 cm height) of a plastic tube (25 cm diameter) was used to acquire a static dorsal thermal image. IR images were taken immediately before and after exercise tests, namely at resting and recovery (Fig.2G), respectively. IR images were analyzed with FLiR Tools software (Flir, Boston)^1,25^.

### Statistics

Data are presented as mean ± Standard Error of the Mean (SEM). Data were evaluated through Student’s t-test or Analysis of Variance (ANOVA), followed by the Bonferroni post-hoc test. The differences were considered significant when P < 0.05.

### Data Availability

The datasets generated and analyzed during the current study are available from the corresponding author on reasonable request.

## Acknowledgements

The work was supported by Prémio Maratona da Saúde, CAPES-FCT (039/2014), FCT (PTDC/NEU-NMC/4154) and ERDF through Centro 2020 (project CENTRO-01-0145-FEDER-000008:BrainHealth 2020). A.S.A.Jr is a CNPq fellow. We would like to acknowledge Flávio N. F. Reis and Frederico C. Pereira (IBILI – Institute for Biomedical Imaging and Life Sciences, University of Coimbra) for making available the treadmill and gas analyzer.

